# Investigating Quinclorac Binding to Human Serum Albumin using Spectroscopy and Molecular Docking

**DOI:** 10.64898/2026.01.08.698511

**Authors:** Le Nguyen, L. Hicks, L. Jodoin, S. Carby, D.S. Sanford, D. Benavides, A. Mishra, R. Yadav

## Abstract

Human serum albumin (HSA) is the most abundant plasma protein and serves as a major transporter and reservoir for a wide range of endogenous ligands, including fatty acids, heme, bilirubin, prostaglandins, and metal ions, as well as numerous exogenous compounds, such as pharmaceuticals. HSA is a 66 kDa monomeric, multidomain protein containing two principal ligand-binding sites, Sudlow sites I and II. Sudlow site I contains Trp214, the sole tryptophan residue in HSA, whose intrinsic fluorescence provides a sensitive probe for studying ligand–protein interactions.

In this study, the interaction between HSA and the persistent organochlorine herbicide quinclorac (QNC) was investigated using steady-state fluorescence spectroscopy and molecular docking. The binding of QNC to HSA is characterized by concentration-dependent quenching of HSA’s intrinsic tryptophan fluorescence. Quantitative analysis revealed a high affinity of QNC for HSA, with ∼ 1 μM dissociation constant (*K*_d_). Temperature-dependent fluorescence quenching experiments indicate a static quenching mechanism. Moreover, a competitive displacement assay using site-specific markers identified Sudlow site I as the primary binding site for QNC in HSA. Induced-fit docking further demonstrated that QNC is accommodated and stabilized by polar interactions, π-stacking, and/or hydrophobic interactions within the Sudlow I binding pocket. Together, these results provide a biochemical and molecular understanding of how QNC, a persistent pesticide, binds to HSA.

## Introduction

Pesticides are environmental pollutants comprising more than 6,000 chemicals currently used to control unwanted organisms, including insects, fungi, and weeds (Carvalho 2017). These environmental pollutants adversely affect human health by accumulating in tissues, causing both acute and chronic toxicity. The widespread and excessive use of these compounds over the past fifty years poses serious health risks for humans, linking exposure to the onset of cancers such as non-Hodgkin’s lymphoma, leukemia, and other solid tumors (Dich *et al*. 1997, Parrón *et al*. 2014, Luo *et al*., 2016).

The quinolinecarboxylic acid quinclorac (3,7-dichloro-8-quinolinecarboxylic acid) belongs to a class of highly selective auxin herbicides and is commonly employed to manage weeds in rice cultivation and to control glyphosate-resistant monocots (Coltro *et al*. 2017, Reichert *et al*. 2023). It is also an active ingredient in several popular professional and consumer weed killers, including Drive XLR8, Q4 Plus, and Roundup for Lawns. Its toxic effects on fish and amphibians are pronounced. A recent study has shown that exposure of the Zebrafish liver cell line under in vitro conditions induces hepatotoxicity and cytotoxicity via oxidative stress (Ferrer *et al*. 2024). However, the U.S. Environmental Protection Agency reports that QNC exhibits low acute toxicity via the oral, inhalation, and dermal routes. However, prolonged and repeated exposure in mammals leads to increased body, liver, and kidney weight, pancreatic acinar cell hyperplasia, and adenomas (Environmental Protection Agency, 40 CFR Part 180). Additionally, QNC has an XLogP3-AA value of 3, indicating moderate lipophilicity and hydrophilicity, as well as high bioavailability.

HSA, a promiscuous protein receptor, is the most abundant plasma protein in humans and plays a crucial role in maintaining osmotic pressure between blood and tissues. In blood plasma, it can also bind and transport various endogenous ligands, including metal cations, fatty acids, hormones, vitamins, and toxic metabolites. HSA can also reversibly bind a wide variety of drugs, affecting their absorption, distribution, metabolism, and elimination (Ghuman *et al*. 2005, Fanali *et al*. 2012). The structure of HSA is characterized by three alpha-helical domains that provide multiple binding sites, which may exhibit cooperativity. These alpha-helical domains, numbered I, II, and III, are each divided into two subdomains, A and B (Fig. 1). Each of these sub-domains has one or two specific binding sites that account for different ligand-binding properties and interact with hydrophobic molecules, long chain fatty acids, bilirubin, charged drugs like warfarin, aspirin, diazepam, ibuprofen, triiodobenzoate and many others (He and Carter 1992, Carter and Ho, 1994). Sudlow *et al*. (1975, 1976) and Sjöholm *et al*. (1979) proposed high-affinity principal binding sites (site I and site II) on HSA located in the hydrophobic subdomains IIA and IIIA. The Sudlow I binding pocket is lined by a mix of hydrophobic residues (e.g., Trp214, Leu219, Phe223, Leu234, Leu238, Leu260, Ile264, Ile290, Ala291) and polar residues (Arg222, Arg257, His242) (He and Carter 1992). In contrast, the Sudlow II binding pocket is mostly lined by nonpolar aromatic residues (Ghuman *et al*. 2005). Most ligands bind to HSA in a site-specific manner, with some classified as site I ligands, such as warfarin, oxyphenbutazone, and dansyl-L-lysine (DNS-Lys), or as site II ligands, including ibuprofen and diazepam (Sudlow *et al*. 1975, Ghuman *et al*. 2005).

**Figure 1.**
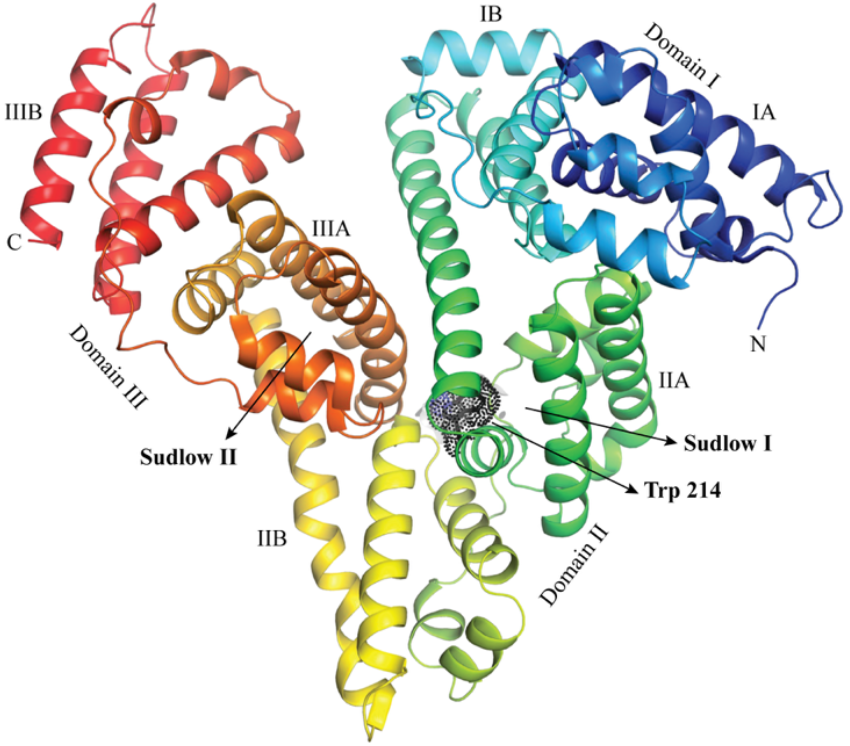
Structure of HSA (PDB ID 2BX8). Three domains and subdomains of HSA have been labeled, and the location of major binding sites (Sudlow I and II) is labeled. The only tryptophan (Trp214) in subdomain IIA is shown as a stick model.

Owing to its remarkable capacity to bind a diverse array of ligands with high affinity, HSA has long been extensively studied to understand its role in drug pharmacokinetics in the pharmaceutical industry. Studies have shown that binding of specific ligands,such as drugs and small molecules, can influence the conformational dynamics, folding pathways, tertiary structure, and packing, thereby impacting the stability of HSA (Chauhan *et al*. 2025, Rogóż *et al*. 2025). Moreover, binding of naproxen and bromocresol green has been shown to alter the stability of HSA’s folded state, highlighting the ligand’s impact on folding and unfolding behavior (Eskew *et al*. 2021). Misfolding of human serum albumin (HSA) has also been associated with diseases such as Alzheimer’s disease and bovine spongiform encephalopathy (Tsao and Meyer 2022).

In this study, we have investigated the interaction and identified the binding site of QNC on HSA using fluorescence spectroscopy, competition assay, and molecular docking. A previous study by Han *et al*. (2009) examined how QNC interacts with bovine serum albumin (BSA). The study suggested weaker interaction and dynamic quenching in BSA upon QNC binding. In contrast, HSA-QNC interaction is characterized by sub-micromolar affinity and a static quenching mechanism.

## Materials and Methods

### Materials

Human serum albumin (≥99%, globulin-free) and QNC (99.7%, PESTANAL® grade) were obtained from Sigma-Aldrich. Sodium phosphate monobasic, sodium phosphate dibasic, and sodium chloride were obtained from Fisher Scientific™. For the site marker displacement assay, Ibuprofen was purchased from MP Biomedicals, and Warfarin and Dansylamide were purchased from TCI Chemicals.

### HSA and Ligand Preparation

A stock of HSA (150-200 μM concentration) was prepared by resuspending the lyophilized powder of HSA in sodium phosphate buffer (20 mM sodium phosphate, 50 mM NaCl, pH 7.4). The concentration of HSA was determined by measuring absorbance at 280 nm with a UV-vis spectrophotometer (Shimadzu UV-2600), using a molar extinction coefficient of 34445 M^-1^ cm^-1^ at this wavelength. A stock solution of 2-20 mM QNC was prepared in DMSO because QNC is insoluble in aqueous buffer. However, the total DMSO percentage was kept below 0.6% during measurement.

### Fluorescence Spectroscopy

Fluorescence measurements were conducted on a JASCO FP-8500 Spectrofluorometer at 25°C by exciting the samples at 295 nm (λ_max_) and recording the emission spectra from 300 to 400 nm. The samples were scanned twice at a scan speed of 50 nm/min, with spectral slits of 2.5 nm for both excitation and emission. Fluorescence quenching of intrinsic tryptophan in HSA was measured by titrating 0.5-25 μM QNC into 5 μM HSA in a 2 mL quartz cuvette. Each spectrum is the average of two scans. All fluorescence spectra were collected across three independent experimental runs. The fluorescence intensity of HSA upon QNC binding was corrected for the inner filter effect by measuring the absorbance of QNC for each ligand concentration using the same measurement setting as with the protein sample. Fluorescence intensity was corrected using the equation (van de Weert and Lorenzo, 2011; Lakowicz, 2006).

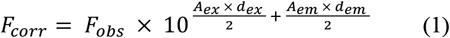

In Eq. 1, *F*_obs_ and *F*_corr_ are the measured and corrected fluorescence intensities of HSA, respectively. *A*_ex_ and *A*_em_ represent the differences in absorbance at the excitation (295 nm) and emission (340 nm) wavelengths, respectively. *d*_ex_ and *d*_em_ are the cuvette pathlengths in the excitation and emission directions (in cm), respectively.

The binding affinity (*K*_d_) for the HSA-QNC interaction was calculated by analyzing the fluorescence quenching data according to the equation (Bakar and Feroz, 2019):

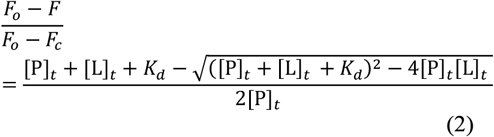

In Eq. 2, *F* is the measured fluorescence, *F*_o_ is the fluorescence in the absence of the ligand, *F*_c_ is the fluorescence of the fully saturated HSA protein at 340 nm, [P]_t_ is the total HSA concentration, and [L]_t_ is the total QNC concentration. The left-hand term in Eq. 2 is equal to the fraction of protein bound. All fluorescence spectra were collected across three independent experimental runs, and the mean and standard deviation were calculated and plotted.

The quenching mechanism upon QNC binding to HSA was investigated by collecting fluorescence titration data at three temperatures (298, 308, and 318 K) and analyzing the quenching results using the Stern-Volmer equation (Lakowicz, 2006).

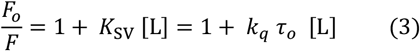

In Eq. 3, *F*_o_ and *F* are the fluorescence intensities of HSA observed in the absence and presence of QNC at 340 nm, respectively; *K*_SV_ is the Stern-Volmer quenching constant, *k*_q_ is the bimolecular quenching rate constant limited by the rate of diffusion of molecules, and τ_o_ is the excited state lifetime in the absence of quencher. To determine *K*_SV_, the data were fitted to Eq. 3 using least-squares fitting. The slope of the plot of Fo/F versus QNC equals the static quenching constant, *K*_SV_.

### Site Marker Displacement Assay

Most small molecules bind to HSA at either Sudlow’s site I (subdomain IIA) or Sudlow’s site II (subdomain IIIA). To identify the binding site of QNC on HSA, a competitive displacement assay was performed using warfarin and ibuprofen as site-specific markers. To determine the binding site of QNC, a complex of HSA (5 μM) and warfarin (6 μM) or ibuprofen (6 μM) was prepared, and fluorescence spectra were collected. To this, an increasing concentration of QNC (5-35 μM or 0-7 molar ratio) was added. The samples were excited at 295 nm, and the emission was recorded over 300-450 nm. The emission signals of warfarin and ibuprofen were analyzed at 379 and 340 nm, respectively.

In addition to warfarin, Dansylamide (DNSA), a fluorescent probe specific for Sudlow’s Site I, was also used to assess QNC binding to HSA (Cheng Er *et al*. 2013). Fluorescence spectra were collected for samples containing 10 μM DNSA alone and 5 μM HSA in the presence of 10 μM DNSA. QNC was then titrated into the HSA-DNSA complex over a concentration range of 5-35 μM (QNC-HSA molar ratio of 1-7). Emission spectra were collected from 400 to 600 nm following excitation at 330 nm.

The percentage of warfarin or ibuprofen displaced by QNC is calculated according to Eq. 4 as follows

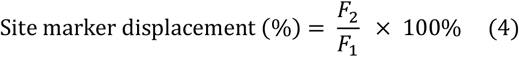

In Eq. 4, *F*_1_ and *F*_2_ are the fluorescence intensities of HSA-bound warfarin, ibuprofen, or DNSA measured in the absence and presence of QNC.

### Molecular Docking

To investigate the binding site of QNC on HSA, molecular docking was performed using the Induced-Fit docking (IFD) module of the Schrödinger Maestro suite (Schrödinger, LLC, New York, NY, 2021). The X-ray crystal structure of HSA with heme and myristic acid (PDB 1N5U) was retrieved from the RCSB PDB database, and atoms of chain A were preserved while associated ligands were removed for docking. The HAS polypeptide chain was optimized and prepared for docking using the Maestro Protein Preparation Wizard, which involved assessing bond order, adding missing hydrogens, and then performing energy minimization with the OPLS3 force field. Water molecules were removed from the protein. Subsequently, the Maestro Receptor Grid Generation module was used to define a 20 × 20 × 20 Å grid centered on the Sudlow I ligand-binding site that contains Trp214.

Ligand QNC was prepared using the Maestro 3D Build module. Then, the Maestro LigPrep module was used to generate QNC conformers, followed by energy minimization using the OPLS3 force field. The IFD workflow combines Glide for initial ligand docking with Prime for protein structure refinement, allowing key side chains and backbone regions within the binding site (selected grid box) to adapt to the ligand. In the first stage, ligands were docked into a softened receptor to generate 5000 initial binding poses; these poses then guided protein refinement, in which residues within 4 Å of the ligand were optimized to accommodate it. Finally, the ligand is redocked into the refined protein structures using high-precision scoring to rank binding modes. Of the total initial poses, the top 400 were selected based on their pose scores and subsequently subjected to energy minimization using the OPLS3 force field. Finally, the top ten poses were generated and ranked based on Glide score, which approximates binding energy by considering receptor-ligand complex energies (Farid *et al*. 2006 and Sherman *et al*. 2006). The highest-ranked docked complex was examined and visualized using Schrödinger Visualizer and PyMOL (Schrödinger, LLC, 2023).

## Results and Discussion

### Fluorescence Quenching Measurements

We have investigated the binding of QNC to HSA by fluorescence quenching. HSA has a single tryptophan residue, Trp214, in the Sudlow I binding site. Selective excitation (λ_ex_ 295 nm) of tryptophan results in fluorescence emission spectra with an intensity maximum near 340 nm in the absence of ligand, which is typical for HSA. Upon addition of increasing concentrations of QNC to HSA, a significant fluorescence quenching was observed without any red shift in the emission maxima at the saturating concentration of QNC (Fig 2A). However, a redshift was observed when a substantially higher concentration of QNC (∼4-5 ligand/protein molar ratio) was added, indicating a structural change in HSA that occurs only at higher QNC concentrations. The ligand QNC absorbs in the UV range around 279, 319, and 332 nm (data not shown). Although the QNC absorbance was not significant at the fluorophore excitation and emission wavelengths, the fluorescence signal from the HSA-QNC interaction was corrected for the inner-filter effect using Eq. 1. A comparison of observed and corrected fluorescence intensity (data not shown) also indicated minimal effect of ligand absorption on the fluorescence intensity of the fluorophore.

**Figure 2.**
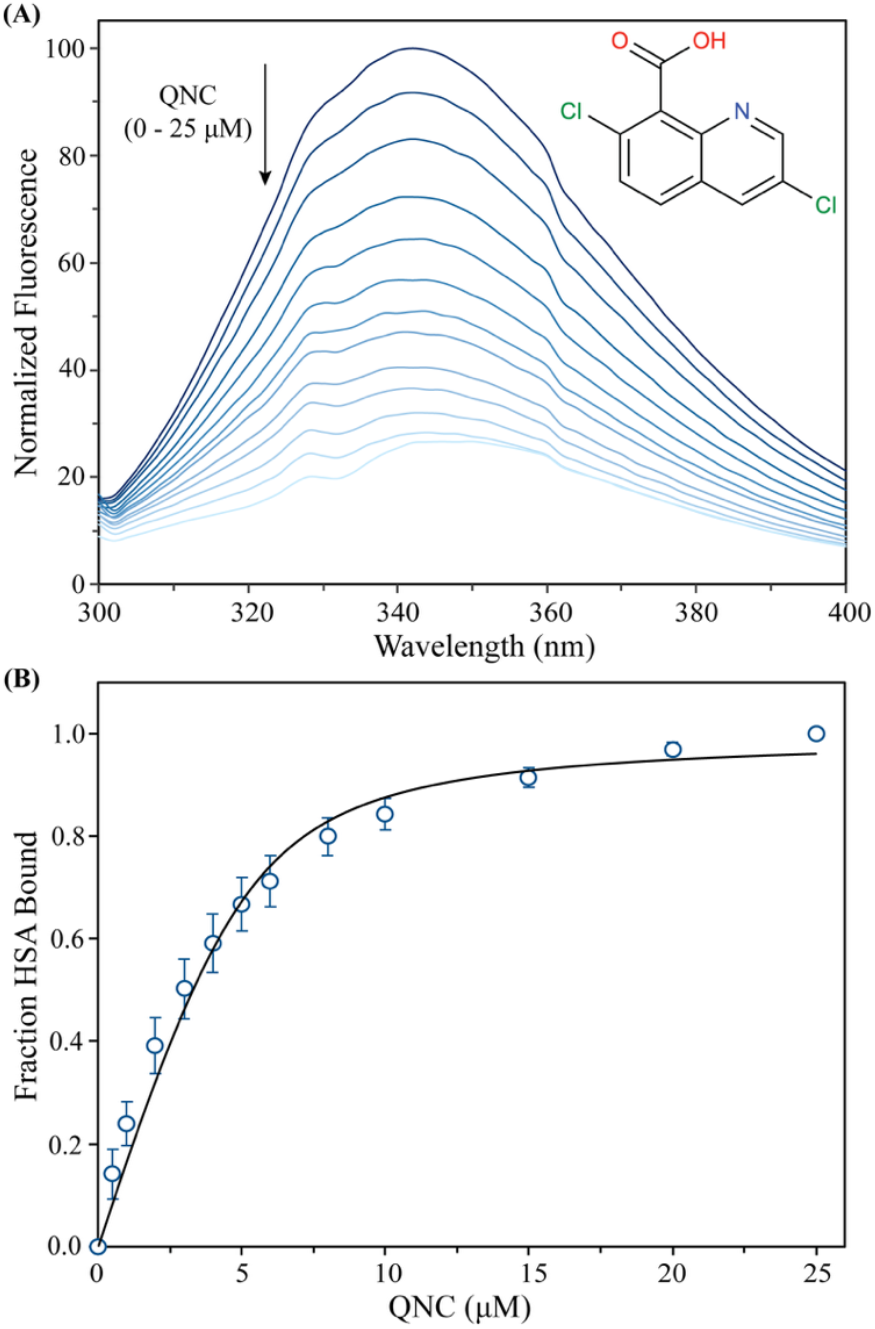
Quenching of HSA tryptophan fluorescence upon addition of QNC. **(A)** Emission spectra (λ_ex_ 295 nm) of 5 μM HSA in the presence of various concentrations of QNC (0.5, 1, 2, 3, 4, 5, 6, 8, 10, 15, 20, and 25 μM) at 298 K. Chemical structure of QNC ligand is shown in the inset. **(B)** The binding isotherm for the interaction of QNC with HSA was obtained from fluorescence quenching measurements. The fraction of HSA bound was calculated from normalized fluorescence changes at the emission maximum (λ_max_ 340 nm) and plotted as a function of QNC concentration. Data points represent the mean ± SD of three independent experiments. The solid curve represents the best fit to a one-site quadratic binding equation (Eq. 2).

Next, we analyzed the fluorescence quenching data above to determine the apparent dissociation constant (*K*_d_) for the HSA-QNC interaction. Although various equations have been employed in the literature to assess the binding affinity of a ligand for HSA, the limitations of these methods have often been overlooked. The challenge in analyzing quenching data stems from the presence of multiple ligand-binding sites in HSA and from distinguishing fluorescence from the bound and unbound fluorophore.

Various methods, rationales, and pitfalls have been reviewed by van de Weert and Stella (2011), Genovese *et al*. (2021), and Bakar and Feroz (2019). Here, we have used the quadratic equation (Eq. 2) to fit the data, which accounts for ligand depletion and quenching that appear to diminish at higher ligand concentrations, thereby reaching a plateau. Fluorescence quenching in HSA upon ligand addition was plotted against ligand concentration (Fig. 2B). Moreover, apparent binding affinity (*K*_d_) was determined by fitting the quenching data to a quadratic equation (Eq. 2), which yielded an apparent *K*_d_ ≈ 1 μM for the HSA-QNC interaction, indicating high affinity of QNC for HSA. During data fitting, we assumed a 1:1 protein-ligand complex and nonzero fluorescence for the complex. Although these assumptions are not ideal for HSA, which is known to contain 4-5 potential binding sites, we believe they are reasonable given the system’s limitations. This is supported by the near-complete quenching of the fluorescence signal at higher ligand concentrations and the results of the ligand site marker displacement assay with Sudlow I site markers, warfarin, and DNSA (discussed later). Nevertheless, the extent of fluorescence quenching does suggest a strong affinity of QNC for HSA.

### Quenching Mechanism

Quenching can occur through two mechanisms, generally classified as dynamic and static quenching. There is also a third mechanism involving components of both static and dynamic quenching. Dynamic and static quenching can be distinguished by examining the temperature dependence of quenching and fluorescence-lifetime measurements. In dynamic or collisional quenching, an increase in temperature enhances molecular diffusion, leading to more frequent collisions and thus a higher quenching constant. Conversely, in static quenching, increasing temperature destabilizes the quencher–fluorophore complex, which reduces the quenching constant.

A plot of *F*_o_/*F* versus ligand concentration was generated, and the Stern–Volmer quenching constant (*K*_SV_) for QNC binding to HSA was calculated at three temperatures using linear regression analysis of Eq. 3. Stern-Volmer plots (Fig. 3) show good linearity up to a ligand/protein molar ratio of 2, suggesting a single quenching mechanism (static or dynamic) for quencher and fluorophore interaction. Moreover, as the temperature increases, the slope of the Stern-Volmer plot, *K*_SV_, decreases, suggesting a static quenching mechanism, which is characterized by ground-state complex formation between the ligand molecule and the protein (Lakowicz, 2006). It should be noted that beyond a ligand/protein molar ratio of 2, the Stern-Volmer plot shows downward curvature or negative deviation (data not shown), suggesting that the classical Stern-Volmer interpretation can be applied to the data. A downward curvature could result from heterogeneity in the fluorophore’s environment and/or from multiple ligand-binding sites, both of which are likely, given multiple ligand-binding sites in HSA. More extensive experiments are required to investigate QNC’s propensity to bind other ligand-binding sites, such as subdomains IB, IIIA, and IIIB in HSA, which are beyond the scope of the current study.

**Figure 3.**
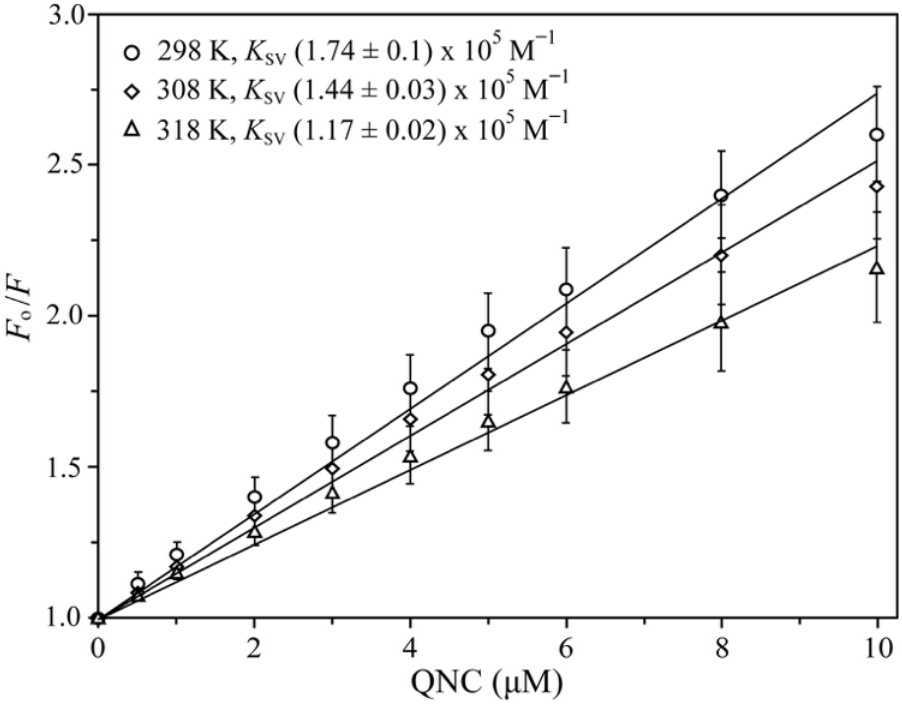
Plot of *F*_o_/*F* against QNC for the calculation of the Stern–Volmer constant (*K*_SV_) (Eq. 3) at three different temperatures. The lines represent linear regression curves forced through the origin.

The binding of QNC to BSA has been studied previously using fluorescence and UV–visible spectroscopy (Han *et al*. 2009). They showed quenching of BSA fluorescence upon QNC binding. However, the extent of quenching was much lower than that observed in HSA upon addition of QNC. Moreover, their study indicated that the Stern-Volmer constant increased with temperature and suggested a dynamic quenching mechanism for BSA fluorescence upon QNC binding. Although BSA and HSA share ∼76% sequence homology, this highlights the significance of structural differences between them.

### Displacement of Site Markers by QNC

We further employed the site marker displacement assay to determine the binding site of QNC on HSA using warfarin and ibuprofen as site markers. The binding sites of these markers on HSA are well characterized and are therefore commonly used to determine the binding sites of small molecules on HSA (Su *et al*. 2021; Qureshi and Javed 2022). Sudlow I in subdomain IIA and Sudlow II in subdomain IIIA are the most prominent ligand-binding sites in HSA, and most ligands show selectivity for one of these sites. While warfarin binds specifically to Sudlow I in subdomain IIA, ibuprofen binds to Sudlow II in subdomain IIIA.

For the site marker displacement assay with warfarin, we first saturated the Sudlow I site in HSA (5 μM) with warfarin (6 μM), resulting in a significantly stronger fluorescence emission at 379 nm than in free HSA. To evaluate QNC’s ability to displace warfarin from HSA, QNC was gradually added to the HSA-warfarin complex until a QNC/HSA molar ratio of 7 was reached. This resulted in a significant, concentration-dependent reduction in the fluorescence emission of HSA-warfarin at 379 nm (Fig. 4A), indicating that QNC was able to displace warfarin from the Sudlow I binding site in HSA. A similar experiment using ibuprofen as a site marker for Sudlow II showed no decrease in the % displacement of ibuprofen by QNC. In contrast, the addition of QNC showed nearly a 50% decrease in warfarin binding (Fig. 4C). This suggests that QNC binds to the Sudlow I site rather than the Sudlow II site.

**Figure 4:**
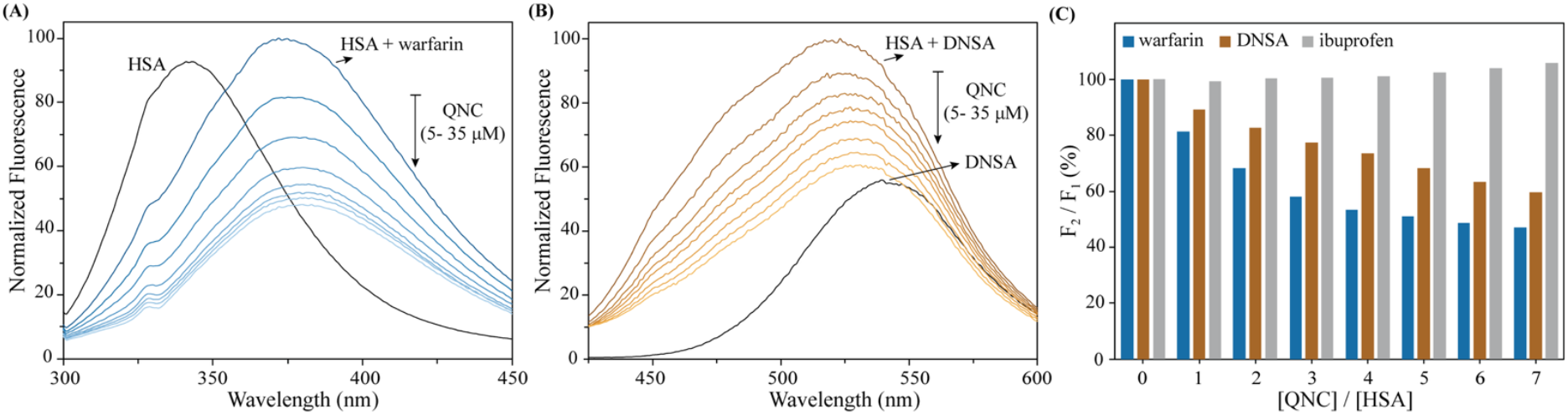
**(A)** Fluorescence emission (λ_ex_ 295 nm) spectrum of warfarin (6 μM) in the presence of HSA (5 μM) and increasing concentrations of QNC (5-35 μM). **(B)** Fluorescence emission (λ_ex_ 330 nm) spectra of DNSA (10 μM) in the absence and presence of HSA (5 μM) and following titration of the HSA-DNSA complex with increasing concentrations of QNC (5-35 μM). Spectra were normalized to facilitate comparison of fluorescence changes. **(C)** Bar graph showing the percent displacement of warfarin, DNSA, and ibuprofen from HSA upon addition of QNC at 298 K. Warfarin and DNSA were used as site markers for specific binding to subdomain IIA, and ibuprofen was used as a marker for subdomain IIIA.

Another competitive displacement assay was performed using DNSA, a site-selective probe for the Sudlow I site of HSA. In contrast to the HSA-warfarin displacement assay, where protein tryptophan was excited at 295 nm, in the HSA-DNSA displacement assay, DNSA was excited at 330 nm. Moreover, DNSA alone exhibited relatively weak fluorescence with an emission maximum near 540 nm. Upon addition of HSA, the fluorescence intensity increased, and the emission maximum shifted to a shorter wavelength (523 nm), indicating the transfer of DNSA from the aqueous phase to the Sudlow I binding pocket of HSA. Subsequent titration of the HSA-DNSA complex with QNC (5-35 µM) resulted in a concentration-dependent decrease in DNSA fluorescence intensity without a substantial change in the emission maximum (Fig. 4B) indicating displacement of DNSA from its binding site. The results indicate that QNC competes with DNSA for binding to HSA and therefore binds at the Sudlow I binding pocket. The displacement assay with warfarin and DNSA indicates that Sudlow I is a major binding site for QNC on HSA. Despite these results, transient or weaker binding of QNC to other ligand-binding sites on HSA cannot be ruled out and requires further studies to ascertain this.

### Molecular Docking

After determining the fluorescence and identifying the binding site of QNC on subdomain IIA in HSA, we employed Schrödinger’s induced fit docking to investigate molecular interactions and determine the amino acid residues involved in QNC binding with HSA. Induced-fit docking allows the selected grid-box residues, including side chains and backbone, to be flexible, if necessary, while keeping the remaining residues on the protein rigid. The initial ligand docking is performed with Glide, followed by structure refinement with Prime to account for side-chain flexibility. By explicitly modeling mutual adaptation of ligand and receptor, the IFD protocol provides more realistic binding poses and improved accuracy.

The best-docked pose of QNC in the Sudlow I binding pocket is shown in Fig. 5. It also suggests the role of Trp214 in stabilizing the QNC in the HSA binding pocket by forming a *π*-*π* interaction with the aromatic ring of tryptophan and the ligand (Fig 5, inset). The binding pocket also displays the amino acid residues in the vicinity of the QNC molecule, highlighting their side chains and key interactions with QNC. For example, positively charged residues Lys195, Lys199, Arg218, and Arg222 are positioned near negatively charged or polar regions of the ligand, indicating electrostatic interactions and hydrogen bonding that stabilize binding. The molecular docking revealed that most of these interactions involved the (deprotonated) carboxylic acid attached to the ligand’s phenyl ring. The docking showed LYS199 interacting with the deprotonated oxygen, while LYS195 interacts with both the deprotonated oxygen and the pi-bonded oxygen. His242 also contributes to polar interactions, while aromatic and hydrophobic residues such as Phe211, Trp214, Phe223, Leu219, Leu238, Ile290,Val293, and Ala residues form a nonpolar environment that supports hydrophobic packing around the ligand scaffold. This suggests that hydrogen bonding, electrostatic interactions, and hydrophobic interactions primarily stabilize the HSA-QNC complex and provide a plausible explanation for QNC’s strong affinity for HSA.

**Figure 5.**
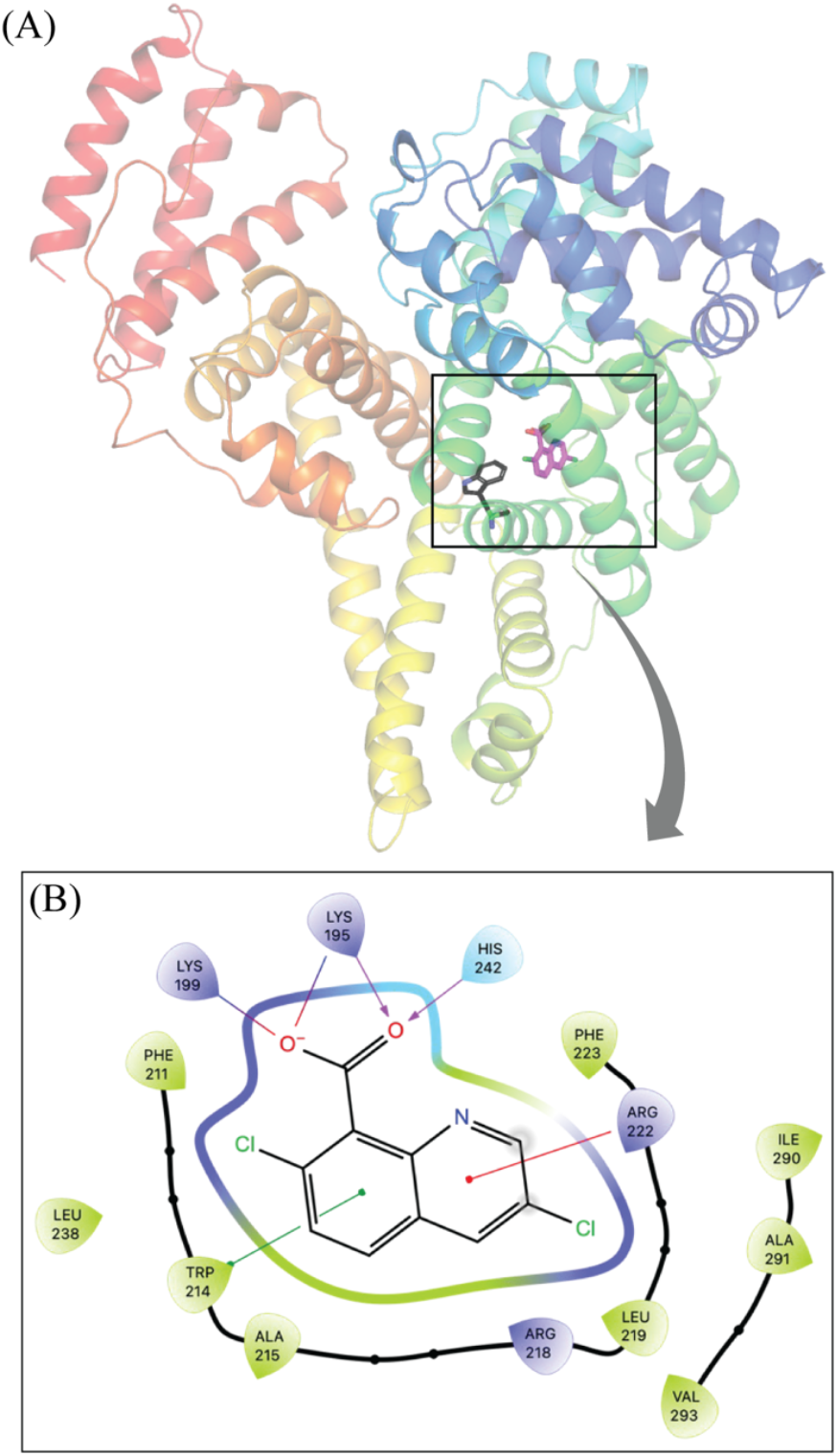
Molecular docking of QNC (magenta) with HSA showing the best docked pose in the Sudlow I binding site. The amino acid residues interacting with QNC are shown (inset). Interaction is stabilized by H-bonding, electrostatic,*π*-cation, and *π*-*π* interaction.

## Conclusion

Fluorescence quenching, site marker displacement, and molecular docking collectively demonstrate that QNC binds strongly and specifically to human serum albumin (HSA). QNC induces pronounced quenching of HSA tryptophan (Trp214) fluorescence without a significant shift in emission maxima at physiologically relevant concentrations, indicating that binding occurs without major perturbation of the local protein environment. Only at higher ligand-to-protein ratios does a red shift appear, suggesting possible structural effects at elevated QNC concentrations. Quantitative analysis of corrected fluorescence quenching data yielded an apparent dissociation constant (*K*_d_) of approximately 1 μM, reflecting high-affinity binding of QNC to HSA. Stern– Volmer analysis showed linear behavior at low ligand/protein ratios and a decrease in the quenching constant with increasing temperature, consistent with a predominantly static quenching mechanism arising from ground-state complex formation between QNC and HSA. Competition with site markers further established that QNC preferentially binds to Sudlow site I (subdomain IIA), as evidenced by significant displacement of warfarin and DNSA but not ibuprofen. Docking analysis revealed that QNC interacts with residues of the Sudlow I binding pocket (notably Lys195, Lys199, Trp214, Arg218, Arg222, and His242).

Overall, the results indicate that QNC forms a stable, high-affinity complex with HSA at Sudlow site I through a combination of electrostatic, hydrogen-bonding, and hydrophobic interactions. This strong binding to HSA is likely to influence the pharmacokinetic behavior of QNC in vivo, underscoring the importance of albumin binding in modulating its distribution and bioavailability.

## Acknowledgements

The University of Arkansas – Fort Smith Physical Sciences Department for Research Resources

